# The accuracy of estimating time to contact in transversal motion and head-on motion

**DOI:** 10.1101/487942

**Authors:** Asieh Daneshi, Hamed Azarnoush, Farzad Towhidkhah, Delphine Bernardin, Jocelyn Faubert

## Abstract

The ability to estimate precisely the time to contact (TTC) of the objects is necessary for planning actions in dynamic environments. TTC estimation is involved in many everyday activities with different levels of difficulty. Although tracking a ball that moves at a constant speed and estimating the time it reaches a non-moving target can be fairly simple, tracking a fast moving car with inconsistent speeds and estimating the time it reaches another car is rather difficult. In this study, we asked participants to estimate TTC of an object in transversal and head-on motions. The object became invisible shortly after movement initiation. The results demonstrated that TTC estimation for transversal motion is more accurate than for head-on motion. We present a mathematical model to explain why humans are better in estimating TTC for transversal motion than for head-on motion.

## 1. Introduction

In everyday life, there are many situations that require us to either avoid or intercept a moving object, even when they are not continuously in view. These objects may be on a collision path with the observers (head-on motion) or not, such as when they pass from one side to the other in front of the observer (lateral motion). Examples of head-on motion include hitting or catching a ball or driving in a street alongside other vehicles, while confronting vehicles when crossing a street is an example of transversal motion. Estimation of time to contact (TTC), which is the time it takes for an object to reach an observer or a particular place, is critical in these situations. During the past decades, several studies have been conducted to understand different aspects of time to contact estimation processes in humans and animals. However, there are still many unanswered questions. One of these questions is what causes the difference between TTC estimation for head-on versus transversal motions. In this study, we first conducted a simple experiment in a 3D environment to examine different aspects of TTC estimation in head-on and transversal motions. We then used a mathematical model to explain the results.

The ratio of an object’s size to its rate of expansion, called tau, is proposed as a prominent source of information for TTC judgments [1-3].

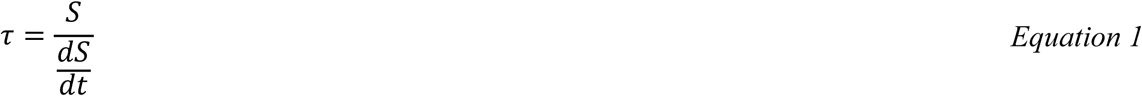

While this provides a reliable judgment of TTC in many situations, it has been shown that a number of additional sources of information are used by the observers, such as luminance of the object, the angular gap between the object’s current position and its final destination, and binocular disparity [4-7].

Different mathematical models were proposed for TTC computing. An extension of the tau formula, termed ‘tau-margin’ was formulated to encompass changes in both angular size (looming), θ, and the angular gap size, ϕ (see Figure 1) [8]. According to this formula, time-to-arrival can be specified as:

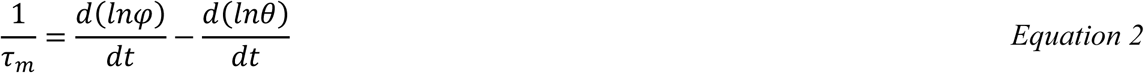

**Figure 1.**
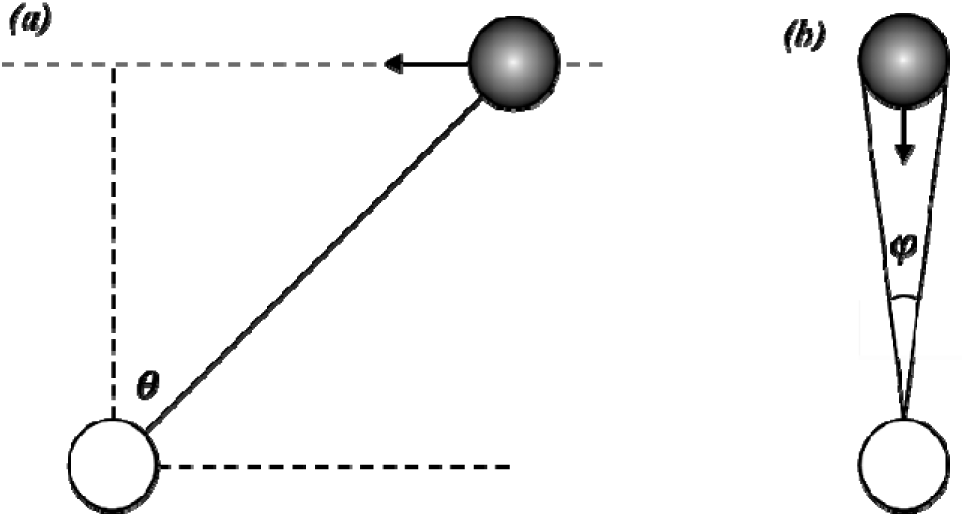
Optical variables computed and used by the model for estimating time-to-contact. (a) Transversal motion, (b) head-on motion. The white balls show the observer, the black balls show the moving object, and the arrows sticking to the black balls show the direction of the black balls ‘ motion.

This is a general equation that simplifies to the TTC condition as proposed by [1] when the object moves on a trajectory 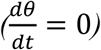, and to the simple 2D gap closure condition when the object does not expand 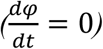. It has been shown that observers are sensitive to the combination of these optical variables (*ϕ* and θ), though with unequal weighting [9]. Also, it was found that observers are sensitive to both the expansion and orientation components of object motion trajectories, including observer self-motion [3, 10].

The formulation of tau is based on a first-order description of object velocity and thus does not consider accelerations. It has been shown that observers perform interceptive actions based on only the linear estimate of tau, even when confronted with accelerating objects [11]. Furthermore, it has been shown that observers are generally poor at accounting for accelerations [12]. A similar result was reported in another TTC estimation study, in which the object moved in depth but not on a collision path to the observer (transversal motion) [13, 14]. Altogether, these results propose that in general, time-to-contact estimations are based on the combination of unambiguous first-order angular velocity estimates.

In 2005, a functional imaging study was conducted to determine the brain areas activated during TTC estimation of head-on and transversal motions (respectively called looming and gap closure tasks in that study). The researchers suggested the presence of separate mechanisms for these tasks [15].

The tasks that are often considered in laboratory-based TTC studies can be classified into two main groups: coincidence anticipation (CA) and relative judgment (RJ) tasks. In CA tasks, participants should make a simple response (e.g. press a button) when the moving object reaches a particular place, called contact point [16]. In an important type of CA tasks, often referred to as prediction motion (PM) tasks, the moving object disappears before reaching the contact point or hides behind a cover. Then, the participant must do a simple action (e.g. press a button) temporally coincident with the time that is expected for the moving object to reach a specific point. The PM paradigm can be used as a relatively straightforward method to assess individuals’ ability to estimate absolute TTC (e.g. [16]). The main purpose of PM tasks is to understand which visual information observers use to estimate TTC. Most PM studies have focused on transversal motion. One of the first studies which has considered head-on motion [17], found that the accuracy of TTC judgments in a PM task decreases as actual TTC increases, and errors typically consist of underestimations. Furthermore, TTC judgments were more accurate in transversal motion than in head-on motion [17]. Later, other researchers obtained similar results when they were studying the contribution of perception and cognition in TTC judgment [18-20]. Some researchers evaluated the parameters that influence contact-time judgments and concluded that the position and speed are important in TTC estimation but they are not the only parameters that individuals use to estimate TTC. A modification of the tau formula, the tau parameter, which is the ratio of visual angle to its derivative through time *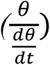,* better estimates the TTC. Recently, several PM studies further supported the notion that an increase in actual TTC decreases the accuracy of TTC judgment [21-23]. However, the main goal of these studies was to study the role of attention in estimating time to contact of one or more objects, and have focused only on the transversal motion from the third person perspective. In 2015, an experiment was conducted to evaluate the contribution of velocity and distance to time estimation during self-initiated time-to-contact judgment [23]. In this experiment, a ball moved horizontally and the participants were asked to launch a stationary ball in a vertical direction towards the first ball, in such a way that they collide. In this study participants had to estimate a kind of non-transversal motion path. However, the experiment was 2-dimensional (2D) and did not represent head-on conditions. Indeed, the participants viewed the experimental space from the top view (2D) as a third person observer. Therefore, whatever the motion trajectories of the two balls they were perpendicular to each other, this could not be categorized as a head-on condition. In our study, we simulated a virtual 3-dimensional environment so that the participants could perceive transversal motion and head-on motion conditions similar to real-world situations.

## 2. Materials and methods

### 2.1. Apparatus

#### a. Subjects

Twenty students from Amirkabir University of Technology-Tehran Polytechnic (10 women, 10 men, age 24.85 years±3.80 (mean±SD), min age 19, max age 31) voluntarily participated in this study. All participants had normal or corrected-to-normal visual acuity. They were healthy and without any known oculomotor abnormalities. Participants were not informed of the experimental hypothesis and gave written informed consent to their participation in the experiment. Iran University of Medical Sciences ethics review board approved all experimental protocols.

#### b. Procedure

Participants sat on a chair facing a 17″ computer display located at a viewing distance of approximately 50 cm, in a room with normal light. Stimuli were generated with Unity3d, and presented on a desktop computer equipped with a 2.90 GHz Intel Corei7 processor. The screen resolution was 1920×1080 pixels (horizontal by vertical) and the display rate was 60 Hz.

The total experiment was divided into two blocks: The Transversal motion experiment and the head-on motion experiment. For each participant, one of these two blocks was selected randomly and after completing that part and taking a short break (five minutes), participants were tested on the remaining block.

### 2.2. Transversal motion experiment

In this experiment, time-to-contact (TTC) estimates for a target car (3 cm length, 1.3 cm width, 1.1 cm height) moving at constant speed in frontoparallel plane from right to left were obtained using a prediction motion (PM) task (see [24]). The constant speeds were randomly selected from three values: 3 cm/s, 4.5 cm/s, 6 cm/s. After 2 seconds, the car became invisible. The point that the car disappeared was the same in all trials, so the initial position of the car was arranged so that the visible time was 2 seconds for all trials (for 3 cm/s, 4.5 cm/s, 6 cm/s the distance between initial position of the target car and disappearance point was respectively 6 cm, 9 cm, 12 cm). To make it possible to study the influence of distance between the observer and the contact point on TTC estimation, observation point was randomly set at a distance of 0 cm, 3 cm, or 6 cm from the red line. Participants were asked to press the spacebar key to start the test. After a delay of 2 s, the car started to move at one of the above mentioned constant speeds, in a horizontal straight line towards the finish line. After 2 s, the car became invisible. The car did not reappear after it became invisible. Participants had 10 seconds to press the “down” arrow key to indicate the instant at which they judged the car would collide with the red finish line. No feedback on TTC estimation error was provided, but a smiley emoji was presented at the end of each trial if the individual finished the trial by pressing the down button and a sad emoji was presented if the subject did not press the down button in the 10-second interval and missed that trial. At the beginning of each trial, we had a countdown from three to one, before the ball started moving. This caused a 2-second delay between two consecutive trials

Figure 2(a) shows a schematic of the transversal paradigm. Nine combinations were generated from three velocity values for the target car and three distance values between the observation point and the red line. Each trial was presented 10 times in random order, for a total of 90 trials. Figure 2(b) shows a screenshot of one of the trials in this experiment and Figure 2(c) shows the schematic of one trial in the transversal paradigm. Values of the parameters in this figure are expressed in Table 1.

**Table 1.**
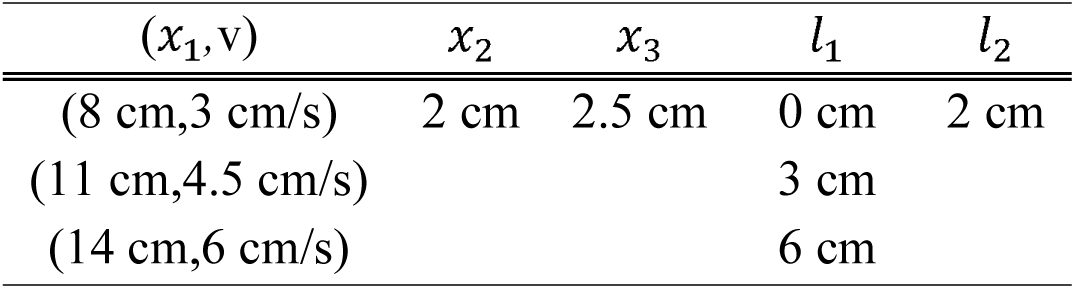
the parameters of the transversal paradigm.

**Figure 2.**
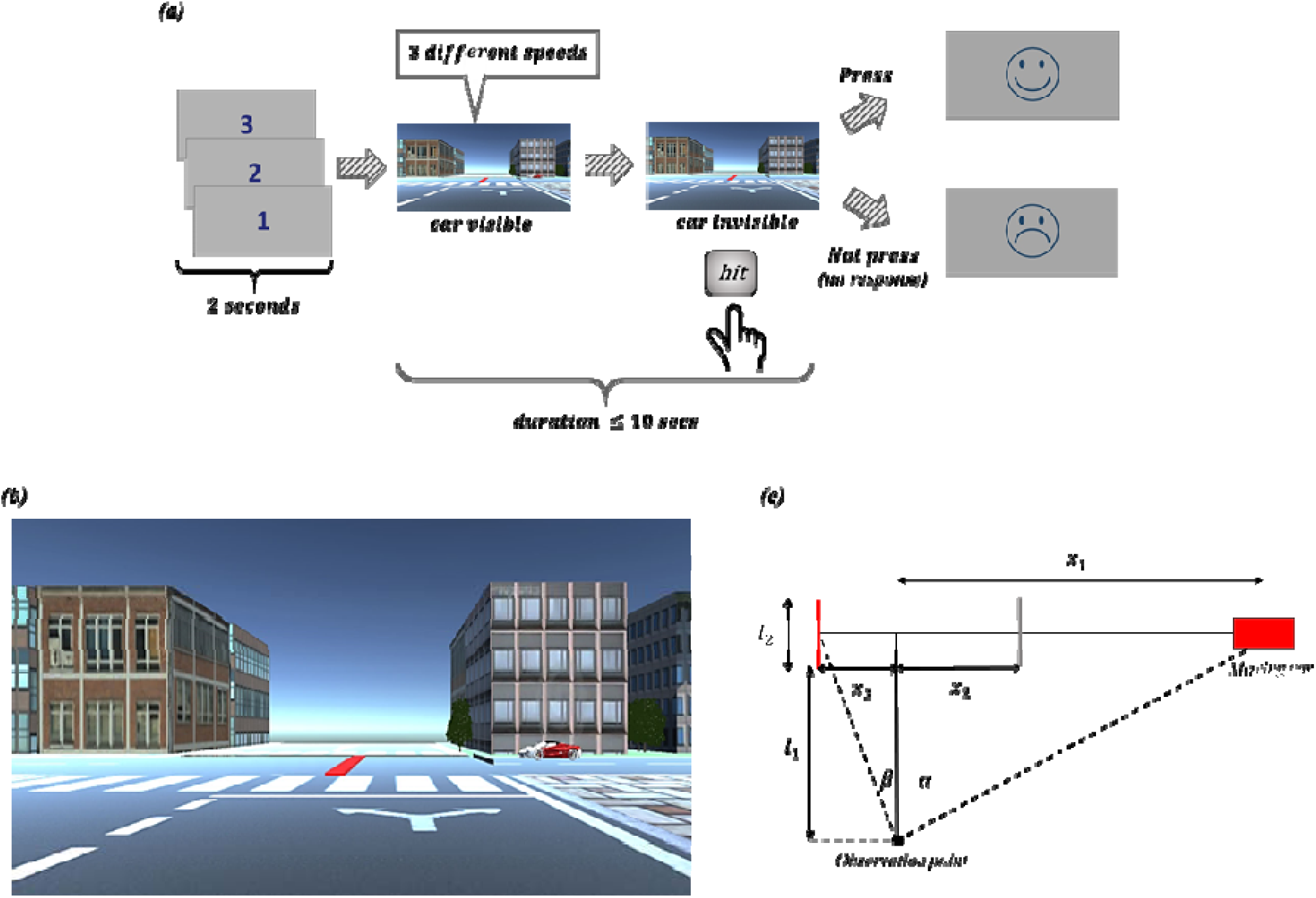
(a) Schematic of the Transversal motion experiment. (b) A screenshot of one of the trials in the transversal paradigm. (c) The schematic of one trial in the transversal paradigm.

### 2.3. head-on motion experiment

In this experiment, all parameters were selected like the Transversal motion experiment, except that the target car moving towards the observation point instead of moving horizontally in frontoparallel plane. Again, no feedback on TTC estimation error was provided, but a smiley emoji was presented at the end of each trial if the individual finished the trial by pressing the down button and a sad emoji was presented if the subject did not press the down button in the 10-second interval and missed that trial. At the beginning of each trial we had a countdown from three to one, before the ball started moving. This caused a 2-second delay between two consecutive trials

Figure 3(a) shows a schematic of the transversal paradigm. Nine combinations were generated from three velocity values for the target car and three distance values between the observation point and the red line. Each trial was presented 10 times in random order, for a total of 90 trials. Figure 3(b) shows a screenshot of one of the trials in this experiment and Figure 3(c) shows the schematic of one trial in the transversal paradigm. Values of the parameters in this figure are expressed in Table 2.

**Table 2.** the parameters of the approach paradigm.

| $l_1$ | $(l_2, v)$ | $l_3$ | $x_1$ |
| --- | --- | --- | --- |
| 0 cm | (6 cm, 3 cm/s) |  |  |
| 3 cm | (9 cm, 4.5 cm/s) | 4.5 cm | 2 cm |
| 6 cm | (12 cm, 6 cm/s) |  |  |

**Figure 3.**
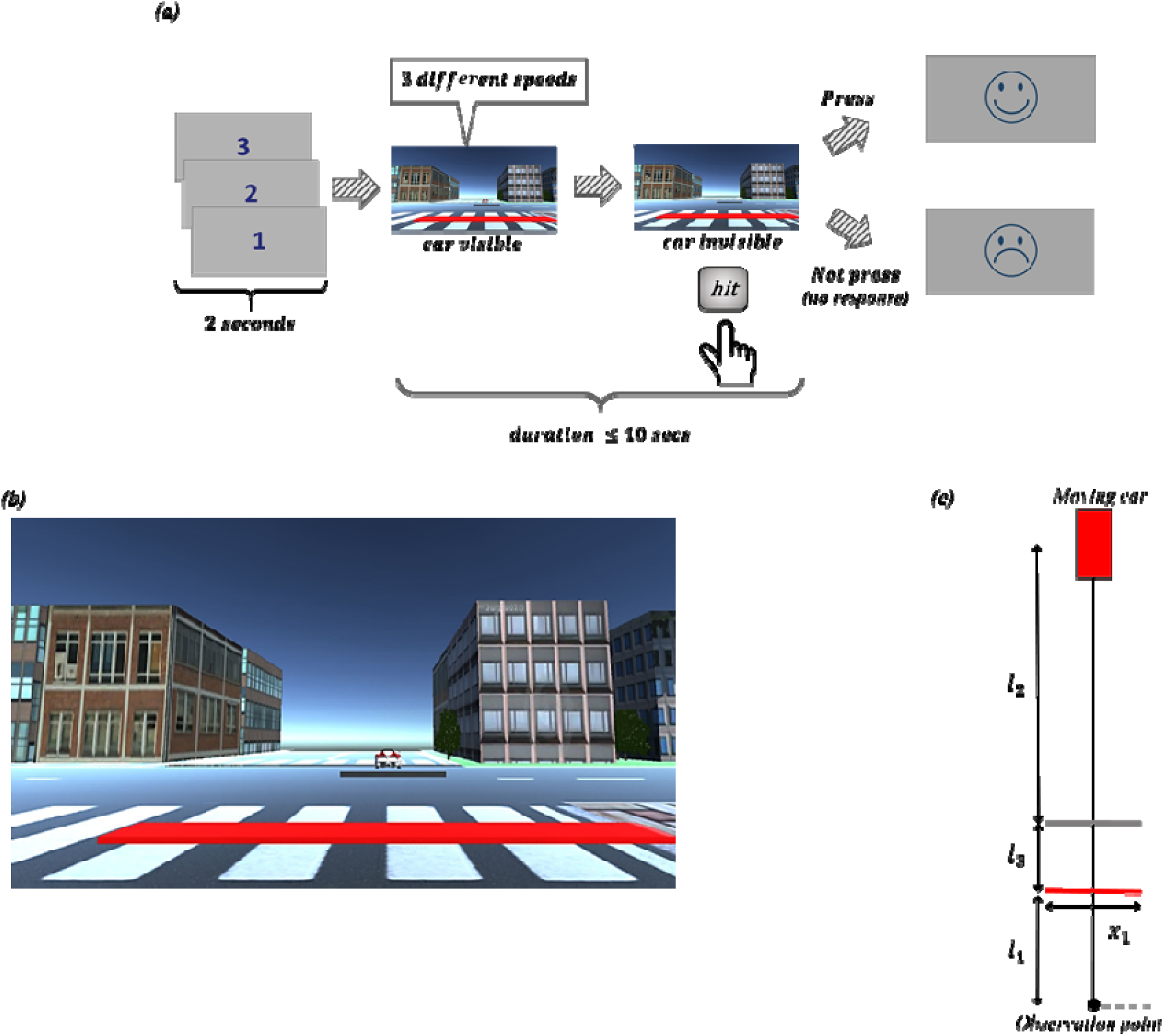
(a) Schematic of the head-on motion experiment. (b) A screenshot of one of the trials in the head-on paradigm. (c) The schematic of one trial in the head-on paradigm.

## 3. Results

First, we studied the estimated response times (the time between the disappearance of the target car and the observer’s response) for the three different target car speeds (3 cm/s, 4.5 cm/s, 6 cm/s). Figure 4(a, b) shows the average estimated response times for all participants and for all trials in these three levels of the target car speed. In both experiments, increasing the target car speed (decreasing the ideal response times) causes a decrease in the average of estimated response times. For the head-on motion experiment, this decrease seems smaller in comparison to the Transversal motion experiment. Figure 4(c, d) shows the average estimated response times for all trials in the three levels of the target car speed. Each dot indicates the average of all response times of one participant in a given target car speed and each color represents one participant. The black plots show the best fit to these points. It is obvious that the slope of the black plot in the head-on motion experiment is lower than the slope of the black plot in the Transversal motion experiment, indicating that the difference between response times for different target car speeds in the head-on motion experiment was smaller than for the Transversal motion experiment. Figure 4(e) shows the average error values for all participants in the three different levels of the target car speed in the Transversal motion experiment (blue bars), and in the head-on motion experiment (red bars).

**Figure 4.**
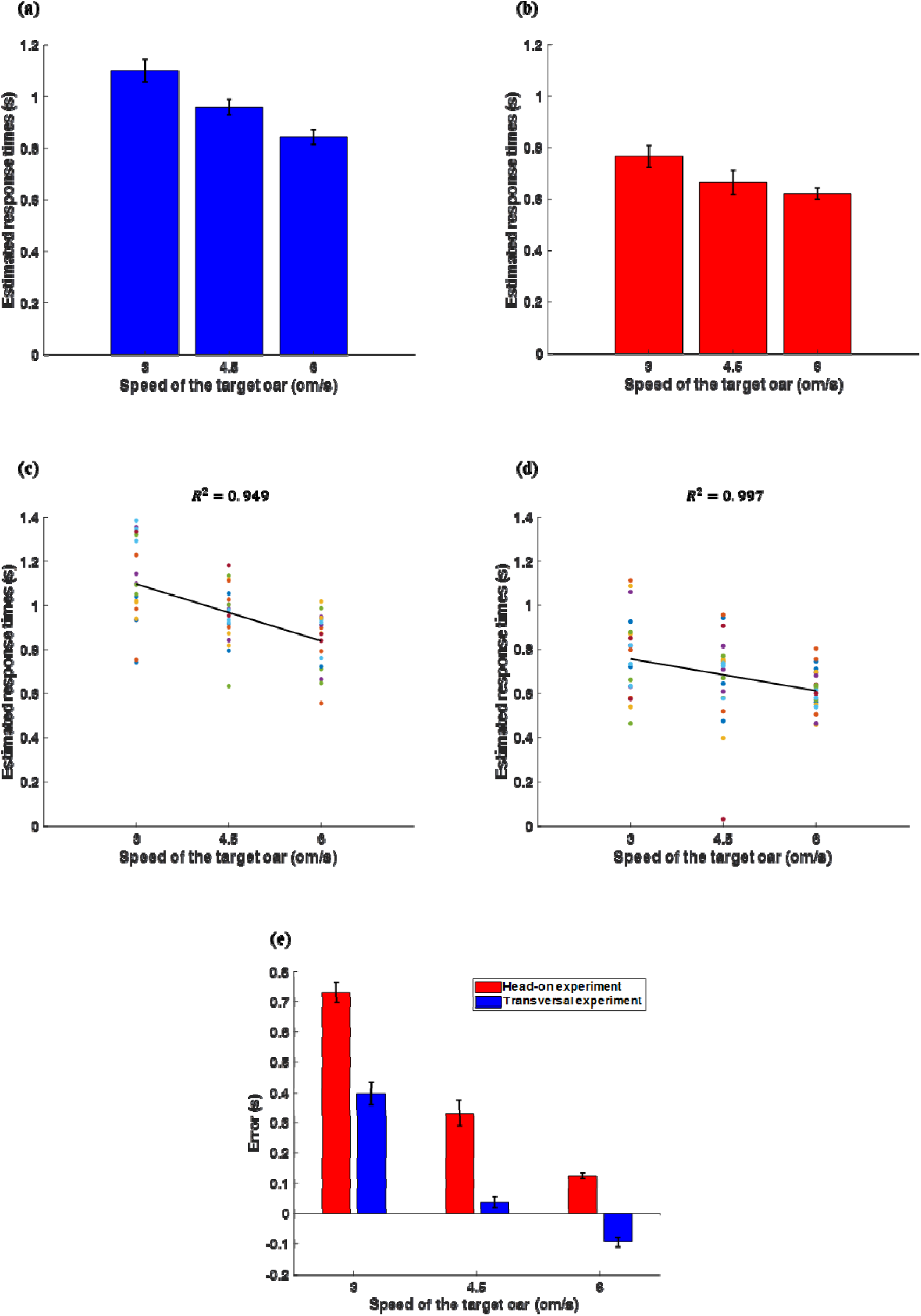
(a) Average response times for the three levels of target car speed for Transversal motion experiment; (b) Average response times for the three levels of target car speed for head-on motion experiment. (c) Estimated TTCs versus target car speeds for the Transversal motion experiment. Each dot indicates the average of all response times of one participant in a given target car speed. Each color is for one participant. Error-bars indicate the standard error of the mean (SEM) for all participants. (d) Estimated TTCs versus target car speeds for the head-on motion experiment. Each dot indicates the average of all response times of one participant in a given target car speed. Each color is for one participant. Error-bars indicate the standard error of the mean (SEM) for all participants. (e) Average TTC error values for the three levels of target car speed, for Transversal motion experiment (Blue bars) and for head-on motion experiment (red bars).

Figure 4(a-d) shows that increasing the target car speed (decreasing the actual response times) have caused a decrease in the estimated response times in both transversal and head-on motion experiments. A one-way ANOVA was conducted on the response times in the Transversal motion experiment and showed a significant difference between the estimated response times in three different target car speeds (F(2,59)=13.965, p<0.001). Then, a post hoc multiple comparisons test showed that the difference between the response times for target car speeds 3 cm/s and 4.5 cm/s was significant (p=0.016), while the difference between the response times for target car speeds 4.5 cm/s and 6 cm/s was not significant (p=0.061). Another one-way ANOVA was conducted on the response times in the head-on motion experiment and showed a significant difference between the estimated response times in three different target car speeds (F(2,59)=3.743, p=0.030). But, a post hoc multiple comparisons test showed that the difference between the response times for target car speeds 3 cm/s and 4.5 cm/s, and also for target car speeds 4.5 cm/s and 6 cm/s was not significant (p=0.201 and p=1.000 respectively).

Also, a one-way ANOVA on the all response times in Transversal motion experiment and head-on motion experiment showed that the difference between estimated response times in these two experiments was significant (p(1,119)<0.001). This can be observed in Figure 4(c,d) that the black plot fitted to response times has different slopes for these experiments (estimated response time = −0.0858 speed + 1.3540 for the Transversal motion experiment and estimated response time = −0.0482 speed + 0.9019 for the head-on motion experiment).

Performance of TTC-estimation in the present study was quantified through calculating error, defined by:

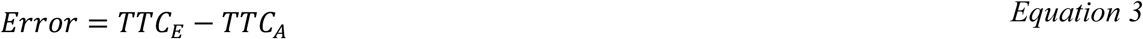

where TTC_E_ (estimated TTC) is the time between the disappearance of the target car and the observer’s response, and TTC_A_ (actual TTC) is the actual time between the disappearance of the target car and when it would have reached the finish line.

Consistent with the pattern of results reported in previous studies [18, 21, 22, 25, 26], accuracy of the responses decreases with larger TTCs (Figure 4(e)). In other words, increasing the actual TTC results in increasing error in TTC estimation.

A three-way ANOVA was conducted on the Error values of all participants to examine the effect of the experiment type, the speed of the target car and the distance between the observation point and contact point on TTC estimation. Experiment types included two groups (transversal and head-on), the speed of the target car consisted of three levels (3 cm/s, 4.5 cm/s, 6 cm/s), and the distance between the observation point and contact point (the point that the target car reaches the finish line) included three levels (0 cm, 3 cm, 6 cm). In all cases, the differences were statistically significant at the 0.05 significance level except for the distance between the observation point and the intersection, meaning that the distance between the observer and the finish line did not make a difference in judging TTC in our experiment. The main difference obtained for paradigm type (Factor 1) yielded an F ratio of F(1,342)=174.812, p<0.001, indicating that the error is significantly higher for approach paradigm (Mean=0.037, SD=0.514) than for lateral paradigm (Mean=-0.419, SD=0.494). The difference obtained for the distance between the observation point and intersection (Factor 2) yielded an F ratio of F(2,342)=0.943, p=0.391, indicating that the effect of this parameter was not statistically significant. Also, the differences obtained for the velocity of the target car (Factor 3) were statistically significant (F(2,342)=662.957, p<0.001; Mean_velocity1_=-0.95, SD_velocity1_=0.482; Mean_velocity2_=-0.919, SD_velocity2_=0.546; Meanvelocity3=-1.020, SD_velocity3_=0.593). The pairwise interaction between parameters was not significant (the interaction between Factor 1 and Factor 2 yielded F(2,342)=1.393, p=0.250, the interaction between Factor 1 and Factor 3 yielded F(2,342)=0.391, p=0.677 and finally the interaction between factor 2 and factor 3 yielded F(4,342)=1.860, p=0.117).

## 1. Discussion

Our results showed that the participants performed significantly better in the transversal motion experiment than in the head-on motion experiment. Here, we present a mathematical model that accounts for why the accuracy of TTC estimation in transversal motion is better than for head-on motion. Consider an object moving laterally from right to left at a constant velocity as shown in Figure 5(a). Points 1 through 5 represent five positions in its trajectory, which are marked at the same distance from each other (the number of the points can be increased arbitrarily, but the present number is enough for the purpose of this paper). The red dot shows the observation point. Tangents of view angles can be written as:

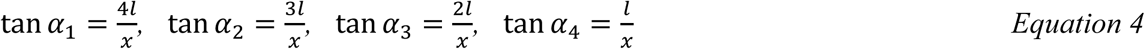

where α_1_, α_2_, α_3_, and a_4_ are visual angles for the points 1 to 4. When the object moves from one of these points to the next, the tangent of its visual angle changes according to the following equations:

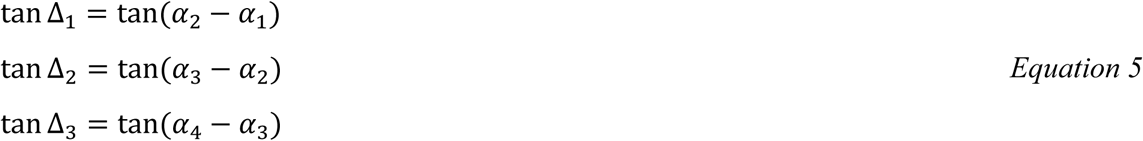

**Figure 5.**
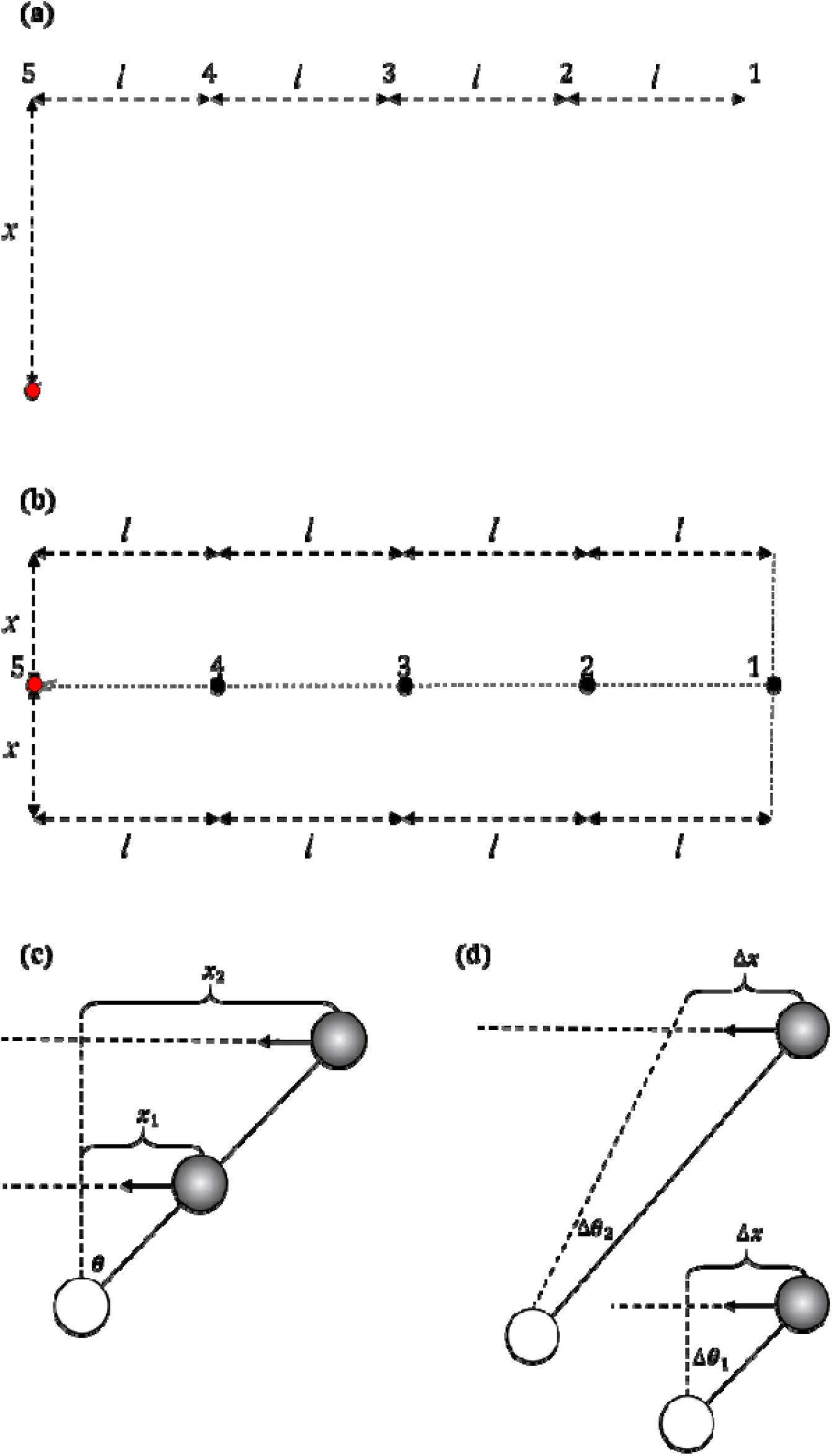
(a) a schematic of five points of transversal motion which are positioned at the same distance from each other. (b) a schematic five points of head-on motion which are positioned at the same distance from each other. (c) a schematic of the transversal motion for two objects moving in parallel. in which the moving object is close to the observer. (d) a schematic of the transversal motion of one object close to the observer and one object far from the observer.

If the distance between two consecutive points is sufficiently small (l very small compared to x), then

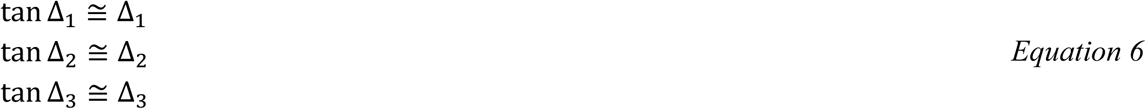

Therefore

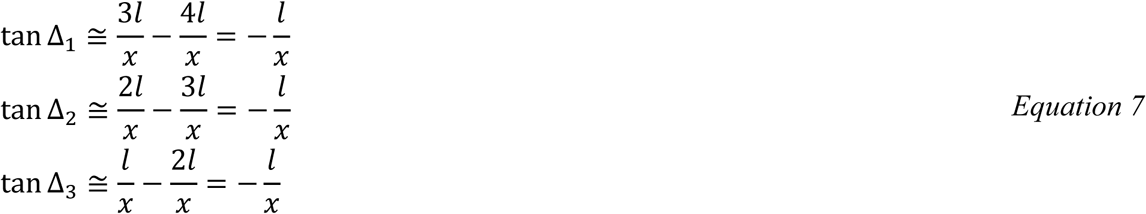

Negativity indicates that the visual angle decreases as the objects moves from Point 1 to Point 5.

Finally,

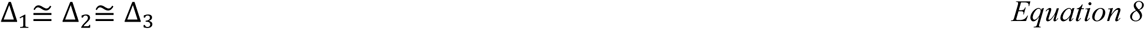

Meaning that visual angle changes at a constant rate, when the object moves laterally on the frontoparallel plane at a constant speed.

Now, consider an object moving towards the viewer. Again five positions of its trajectory, having the same distance from each other are marked (points 1 to 5) and the red dot shows the observation point. Here, tangents of visual angles can be written as:

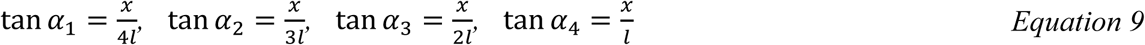

where α_1_, α_2_, α_3_, and α_4_ are visual angles for the points 1 to 4 and ls and × are remarked in Figure 5(b). When the object moves from one of these points to the next, the tangent of its visual angle changes according to the following equations:

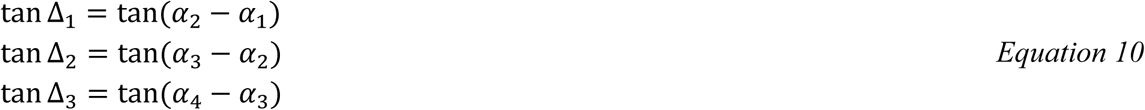

If the visual angles be small enough (x be very small compared to l), then

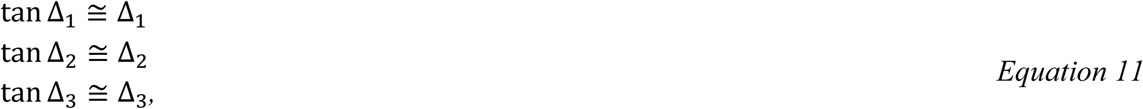

and

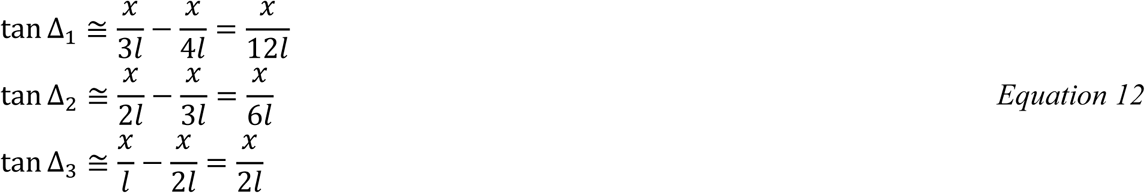

It is obvious that the visual angle increases as the object moves towards the observer.

Therefore,

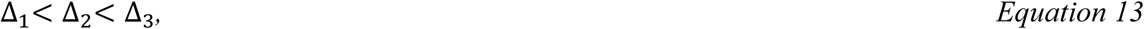

meaning that the rate of change in view angle is not constant when the object approaches the observer or moves away from him/her. As the object moves towards the observer, the angular velocity increases and the observer perceives an accelerated movement, while in the transversal motion, the observer perceives a constant angular velocity. Humans can perceive that an object moves at a non-constant speed, but they are not able to estimate the acceleration value in moving visual stimuli [27, 28]. As a result, the performance for transversal TTC estimation is better than that of head-on TTC estimation.

As mentioned above, a three-way ANOVA was conducted on the Error values of all participants to examine the effect of the experiment type, the speed of the target car and the distance between the observation point and contact point on TTC estimation. This showed that the distance between the observation point and contact point does not have a significant impact on TTC estimation. However, based on the motion parallax theorem we expected a significant impact. According to this theorem, closer objects appear to move faster rather than objects that are farther away [29]. Consider an object moving in parallel trajectories from right to left (gray circles in Figure 5(c,d)), while an observer (white circles in Figure 5(c,d)), tracks their movements. For equal angular velocities, the farther object must move at a higher velocity than the nearer object (Figure 5(c)), because for the same time-interval, the distance that the farther object should travel is larger than the distance that nearer object should travel.

If these objects move at the same velocity, the angular velocity for the closer object is more than the other object (Figure 5(d)). For the same time interval and the same velocity, the angular velocity 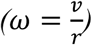 of the nearer object is larger than the angular velocity of the farther object. where *ν* is the velocity, *ω* is the angular velocity, and *r* is the distance between the observer and the moving object. It is obvious that for the farther objects *r* is larger, and at the same ν, *ω* is smaller for the farther object. That is why closer objects appear to move faster than objects that are farther.

But, changing the distance between the observer and the finish line did not cause a statistically significant difference in judging TTC in our experiment. One possible reason for explaining our results is that the values that we selected for the parameters in our experiment do not make a significant difference between the angular velocities; therefore, motion parallax theorem cannot be directly addressed in our experiment.

## 2. Conclusion

We aimed to study the TTC estimation in transversal motion and head-on motion in a 3D environment, similar to the real world. We asked participants to estimate TTC of a target car hitting an end line after it disappeared from sight. Our results showed that the accuracy of TTC estimation for the transversal motion was significantly better than for the head-on motion (Figure 4(e)). We proposed a mathematical model based on behavioral responses for these results. We showed that if an object has a constant speed in the head-on motion, its movement is perceived as accelerated. Previous studies have shown that human observers are not able to estimate the value of acceleration in ordinary situations, and use only the information about time, position, and speed of an object to estimate TTC [30, 31]. This fact together with the fact that human observers perceive constant-speed head-on motions as accelerated, but constant-speed transversal motions without acceleration led us to elucidate why the accuracy of TTC estimation in the transversal motion is better than in the head-on motion.

We also expected that the distance between the observer and the contact point influences the accuracy of TTC estimation. The motion parallax theorem [29, 32-34] can explain that if the speed of two moving objects is the same, the closer object is perceived to move faster than the other object. But, our results did not show a significant effect of this factor. We proposed that this is because, in our experiment, the visual angle for the condition that target car is far from the observer is not much different from the visual angle for the condition that target car is close to the observer. In future studies, experiments can be designed to study these conditions.

Some studies have shown that TTC estimates for each target car speed tends to be biased toward the mean TTC across all target car speeds [24, 25, 35, 36]. We did not observe such pattern in our experiments, maybe because we did not provide participants with feedback on estimation accuracy. Such feedback can result in forming priors for the subjects centered around the mean TTC which would bias estimates toward the mean in each trial. Furthermore, there are other studies that reported overall underestimation of TTCs similar to our results [26, 37].

In summary, we provided a mathematical model based on behavioral results about why the accuracy of TTC estimation for transversal motion is better than for the head-on motion. Future studies should address the influence of the distance between the observation point and the contact point on TTC estimation accuracy. Furthermore, the neural basis for tracking objects in both transversal motion and the head-on motion and the brain areas involved in them needs to be addressed.

## References

[1] D. N. Lee, “A theory of visual control of braking based on information about time-to-collision,” Perception, vol. 5, pp. 437–459, 1976.

[2] H. Hecht and G. Savelsbergh, “Time-to-contact, Advances in Psychology Series,” ed: Elsevier, North Holland, Amsterdam, 2004.

[3] G. A. Geri, R. Gray, and R. Grutzmacher, “Simulating time-to-contact when both target and observer are in motion,” Displays, vol. 31, pp. 59–66, 2010.

[4] R. Gray and D. Regan, “Accuracy of estimating time to collision using binocular and monocular information,” Vision research, vol. 38, pp. 499–512, 1998.

[5] R. Gray and D. Regan, “The use of binocular time-to-contact information,” in Advances in psychology. vol. 135, ed: Elsevier, 2004, pp. 303–325.

[6] S. K. Rushton and J. P. Wann, “Weighted combination of size and disparity: a computational model for timing a ball catch,” Nature neuroscience, vol. 2, p. 186, 1999.

[7] A.-M. Brouwer, J. López-Moliner, E. Brenner, and J. B. Smeets, “Determining whether a ball will land behind or in front of you: Not just a combination of expansion and angular velocity,” Vision Research, vol. 46, pp. 382–391, 2006.

[8] R. J. Bootsma and R. R. Oudejans, “Visual information about time-to-collision between two objects,” Journal of experimental psychology: human perception and performance, vol. 19, p. 1041, 1993.

[9] B. Lorv, “Time-to-Collision of Looming Spherical Objects: Tau Revisited,” 2011.

[10] R. J. Bootsma and C. M. Craig, “Global and local contributions to the optical specification of time to contact: Observer sensitivity to composite tau,” Perception, vol. 31, pp. 901–924, 2002.

[11] D. Lee, D. Young, P. Reddish, S. Lough, and T. Clayton, “Visual timing in hitting an accelerating ball,” The Quarterly Journal of Experimental Psychology, vol. 35, pp. 333–346, 1983.

[12] N. Benguigui, H. Ripoll, and M. P. Broderick, “Time-to-contact estimation of accelerated stimuli is based on first-order information,” Journal of Experimental Psychology: Human Perception and Performance, vol. 29, p. 1083, 2003.

[13] M. K. Kaiser and L. Mowafy, “Optical specification of time-to-passage: observers’ sensitivity to global tau,” Journal of Experimental Psychology: Human Perception and Performance, vol. 19, p. 1028, 1993.

[14] M. K. Kaiser and H. Hecht, “Time-to-passage judgments in nonconstant optical flow fields,” Perception & Psychophysics, vol. 57, pp. 817–825, 1995.

[15] D. T. Field and J. P. Wann, “Perceiving time to collision activates the sensorimotor cortex,” Current Biology, vol. 15, pp. 453–458, 2005.

[16] J. Tresilian, “Perceptual and cognitive processes in time-to-contact estimation: analysis of prediction-motion and relative judgment tasks,” Attention, Perception, & Psychophysics, vol. 57, pp. 231–245, 1995.

[17] W. Schiff and R. Oldak, “Accuracy of judging time to arrival: effects of modality, trajectory, and gender,” Journal of Experimental Psychology: Human Perception and Performance, vol. 16, p. 303, 1990.

[18] J. Tresilian, “Perceptual and cognitive processes in time-to-contact estimation: Analysis of prediction-motion and relative judgment tasks,” Perception & Psychophysics, vol. 57, pp. 231–245, 1995.

[19] S. K. Rushton and P. A. Duke, “Observers cannot accurately estimate the speed of an approaching object in flight,” Vision research, vol. 49, pp. 1919–1928, 2009.

[20] P. R. DeLucia and G. W. Liddell, “Cognitive motion extrapolation and cognitive clocking in prediction motion tasks,” Journal of Experimental Psychology: Human Perception and Performance, vol. 24, p. 901, 1998.

[21] R. Baurès, D. Oberfeld, and H. Hecht, “Judging the contact-times of multiple objects: Evidence for asymmetric interference,” Acta Psychologica, vol. 134, pp. 363–371, 2010.

[22] R. Baurès, D. Oberfeld, and H. Hecht, “Temporal-range estimation of multiple objects: Evidence for an early bottleneck,” Acta psychologica, vol. 137, pp. 76–82, 2011.

[23] Y. Li, L. Mo, and Q. Chen, “Differential contribution of velocity and distance to time estimation during self-initiated time-to-collision judgment,” Neuropsychologia, vol. 73, pp. 35–47, 2015.

[24] W. Schiff and M. L. Detwiler, “Information used in judging impending collision,” Perception, vol. 8, pp. 647–658, 1979.

[25] C.-J. Chang and M. Jazayeri, “Integration of speed and time for estimating time to contact,” Proceedings of the National Academy of Sciences, vol. 115, pp. E2879–E2887, 2018.

[26] J. Steeves, R. Gray, M. Steinbach, and D. Regan, “Accuracy of estimating time to collision using only monocular information in unilaterally enucleated observers and monocularly viewing normal controls,” Vision research, vol. 40, pp. 3783–3789, 2000.

[27] S. N. Watamaniuk and A. Duchon, “The human visual system averages speed information,” Vision research, vol. 32, pp. 931–941, 1992.

[28] A. S. Mueller and B. Timney, “Visual acceleration perception for simple and complex motion patterns,” PloS one, vol. 11, p. e0149413, 2016.

[29] K. L. Norman, Cyberpsychology: An introduction to human-computer interaction: Cambridge university press, 2017.

[30] P. Senot, P. Prévost, and J. McIntyre, “Estimating time to contact and impact velocity when catching an accelerating object with the hand,” Journal of Experimental Psychology: Human Perception and Performance, vol. 29, p. 219, 2003.

[31] A. S. Mueller, “Effects of Motion Pattern Characteristics on the Perception of Visual Acceleration,” 2015.

[32] K. Nakayama, “Biological image motion processing: a review,” Vision research, vol. 25, pp. 625–660, 1985.

[33] P. DeLucia, M. Kaiser, J. Bush, L. Meyer, and B. Sweet, “Information integration in judgements of time to contact,” The Quarterly Journal of Experimental Psychology: Section A, vol. 56, pp. 1165–1189, 2003.

[34] H. Hecht and G. J. Savelsbergh, Time-to-contact vol. 135: Elsevier, 2004.

[35] M. B. Ahrens and M. Sahani, “Observers exploit stochastic models of sensory change to help judge the passage of time,” Current Biology, vol. 21, pp. 200–206, 2011.

[36] M. Jazayeri and M. N. Shadlen, “Temporal context calibrates interval timing,” Nature neuroscience, vol. 13, p. 1020, 2010.

[37] N. Yakimoff, S. Mateeff, W. H. Ehrenstein, and J. Hohnsbein, “Motion extrapolation performance: A linear model approach,” Human factors, vol. 35, pp. 501–510, 1993.

